# Odor-induced persistent neural activity in associative encoding in humans

**DOI:** 10.1101/2024.02.23.581728

**Authors:** Joan Tarrida, Manuel Moreno, Jordi Vidal, David Panyella, Josep Marco-Pallarés, Lluís Fuentemilla

## Abstract

This study explored the impact of brief exposure to odor cues on sustained neural activity during a 6-second delay period before memory encoding of a picture image. Combining univariate and multivariate ERP analytical approaches, our results align with nonhuman data, indicating that odor cues induced sustained neural activity in humans, persisting beyond the odor exposure throughout the nearly 6-second delay period. We observed higher amplitude of sustained ERPs for unpleasant compared to pleasant odor cues. Additionally, participants exhibited more confident memory recall for pictures preceded by unpleasant rather than pleasant odor cues during encoding, underscoring the influence of brief odor cues on memory formation for temporally distant events. In conclusion, this study revealed that brief exposure to odor cues induced sustained neural activity in humans, with distinct effects on memory formation along the pleasantness dimension, emphasizing the lasting impact of olfactory stimuli on cognitive processes.

## Introduction

Odors are integral to our sensory experiences, exerting a robust and long-lasting influence on how we will remember them in the future. For example, when entering the charming cafe, a passing whiff of a nearby overflowing trash bin briefly mars the welcoming ambiance for you. As you settle in and enjoy the coffee, the unpleasant odor dissipates. Yet, every time you reminisce about that cafe visit, the memory of the brief unpleasant scent resurfaces, subtly influencing how you recall the otherwise neutral experience. While this example emphasizes the enduring impact of olfactory experiences in memory formation, little is understood about how the representations after odor offset are generated and used to form associative trace memory.

Persistent neural activity refers to a sustained neural activity increase that long outlasts a stimulus [1, 2]. It is found in a diverse set of brain regions and organisms and several in vitro systems, suggesting that it can be considered a universal form of circuit dynamics that can be used as a mechanism for short-term storage, enabling the brain to link temporally distant inputs into a bound memory trace [3]. Odor-related neural persistence has been seen to be particularly long-lasting [4] and to be a mechanistic conduit supporting the advantage of trace over delay conditioning in animal studies [5]. In humans, smells can also evoke powerful lingering effects on brain activity subsequent to its exposure, especially for unpleasant odor cues [6]. However, while odor induced neural persistence has been identified in the context of associative learning in rodents [7] and in insects [5], this research has been scant at best in humans.

Here, we assessed neural persistence in humans induced by brief exposure to pleasant and unpleasant odors and investigated its impact on memory for upcoming events. The use of pleasant and unpleasant odors is motivated by previous studies in humans describing different responses of the olfactory system depending on their valence [6]. Indeed, studies in nonhuman animals revealed that the olfactory system is particularly suited to detect, and avoid aversive olfactory stimuli [8, 9], and respond specifically in front of unpleasant but not pleasant odors [10]. We measured event-related potentials (ERPs) while healthy adult participants were asked to encode the association between an odor cue and a picture of a scene presented 6 seconds after. ERPs were chosen because, unlike other neuroimaging approaches, they allow tracking neural activity changes leading up to an event, offering fine-grained time resolution. This allowed us to distinguish neural state signals in response to closely occurring events, such as during the encoding of a cue, the offset period, and during the encoding of the upcoming event. We hypothesized that picture images preceded by unpleasant odor cues would be better remembered than those preceded by pleasant odor cues in an online recognition test 24-48 hours after encoding. We also predicted that brief odor exposure would induce a sustained neural activity during the delay period between odor and image, thereby informing about the existence of odor-related neural persistence in humans.

## Materials and Methods

### Participants

Following similar previous studies [11], twenty-six healthy participants (11 women) participated in the study. The range in age was 19 - 34 years old (mean = 24, SD = 4.37). All participants provided informed written consent for the protocol approved by the Ethics Committee of the University of Barcelona. Participants received financial compensation for their participation (20€). One of the participants was excluded due to excessive artifacts on the EEG data during the recording. Two other participants could not complete the recognition test and were excluded from all analysis. Thus, a total of 23 participants were included in the analysis.

### Stimuli

Two types of stimuli were used in this study: odors and pictures. The odors presented were L-Carvone (CAR, diluted in 20% Ethanol) and N-Butanol (BUT, diluted in 30% Ethanol). The odor stimuli were selected based on their expected hedonic properties [12–14]. CAR as pleasant and BUT as unpleasant. Pictures belonged to two different semantic categories: animals and objects (120 animals and 120 objects) used in a previous study [15]. All pictures were iso-luminant. Displayed at the center of a uniform gray background of 33 cd/m^2^ and subtended approximately 17.44 degrees of visual angle at a viewing distance of 70 cm. Iso-luminance was achieved by normalizing the pixel levels integration of each picture to the same intensity of 33 cd/m^2^.

### Experimental Procedure

The experiment consisted of an encoding and a retrieval phase, separated by 48 - 72 hours Figure 1. Task timing and visual stimulus presentation were under the control of Python libraries Pyglet [16] for the encoding task and PsychoPy [17]for the recognition task.

**Figure 1.**
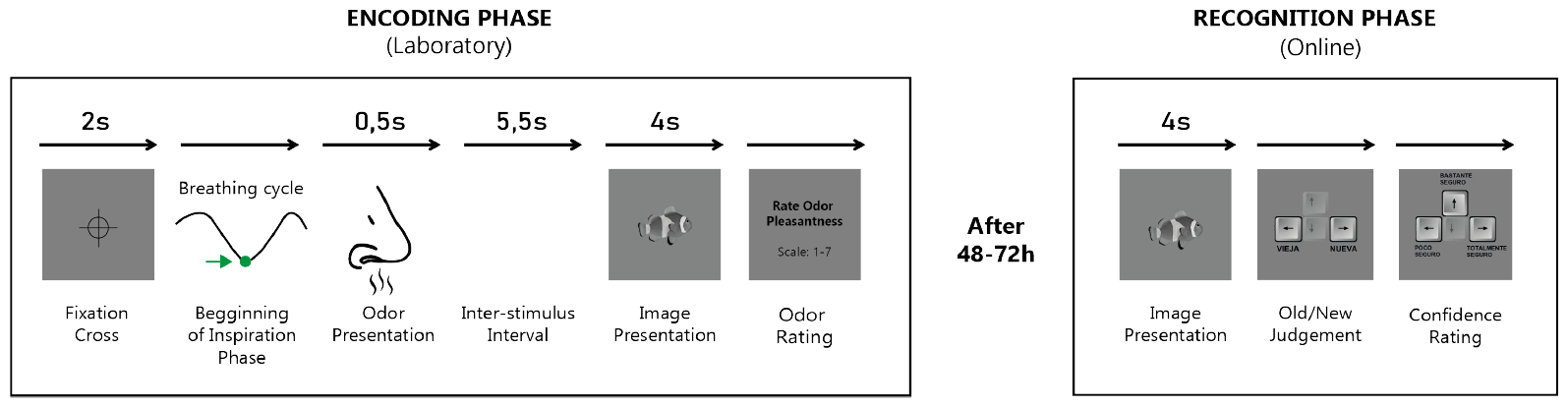
Experimental design. During encoding phase, 120 odor trials were preceding image presentation. The trial began with a fixation cross presented for 2 s. When participant breathing cycle changed from exhalation to inspiration, the odor was sent. After 6 s, a picture of an animal or object was presented on screen during 4 s. The trial ended with participants being asked to rate odor pleasantness in a 1 to 7 Likert scale. As soon as they responded, the fixation cross appeared on the screen. The recognition phase consisted of an online task in which 240 pictures were presented (120 previously presented, 120 new ones). Pictures appeared on screen during 4 s. Subsequently participants were asked to judge the picture as old or new and their level of confidence “Low”, “Medium” or “High.”).

Participants were informed about the two-day structure of the experimental design. On day 1, participants were asked to encode a series of pictures preceded by an odor. Participants were not informed there would be a memory test of the pictures. The encoding phase consisted of 120 trials. In each trial CAR, BUT, and clean air preceded the presentation of a picture. Each type of odor was paired with 40 trials of a specific semantic category. Twenty trials of each category were paired with clean air. For example, in one participant CAR was paired with 40 trials of animals, BUT with 40 trials of objects and clean air with 20 trials of animals and 20 of objects. Odor-semantic category associations were counterbalanced across participants. Clean air was presented continuously during the whole experiment to avoid tactile stimulation when presenting the odors. When odors were presented, a solenoid valve opened for 0.5 s allowing the air to pass through a channel that contained a flask with an odor in a liquid state. When the air touched the surface of the liquid it got odorized.

The trial structure for the encoding phase was as follows. A fixation cross appeared on screen for 2 seconds indicating the beginning of the trial. After these 2 seconds and when the participant breathing cycle, recorded by a respiration belt, changed from expiration to inspiration, a brief odor pulse was sent to the participant nose using an olfactometer. After 6 seconds since the odor has been sent a picture was displayed on screen for 4 seconds. Next, the participant had to rate how much did they like the previous odor. Rating was made in a Likert scale from one to seven, being One very unpleasant, four neutral and seven very pleasant. When the participant rated the odor, the fixation cross appears to the screen again indicating the beginning of a new trial. Each 10 trials there was a pause where participants decided when to continue pressing the space bar.

On day 2, participants performed an online surprise recognition memory test. The 120 pictures of day 1 and 120 new ones were presented. In the recognition test, each trial began with the presentation of one of the 240 pictures. After the 4 seconds, participants had to indicate if the picture was old (i.e., encoded in the previous encoding phase) or new (i.e., not presented during the encoding phase). Subsequently, participants were asked to rate their confidence in their decision, choosing from “Low”, “Medium” or “High.”

### EEG Recording

During the encoding phase, scalp EEG was recorded using a 32-channel system at a sampling rate of 250 Hz, using a BrainAmp amplifier and active electrodes (actiCap) located at 29 standard positions (Fp1/2, Fz, F7/8, F3/4, Fc1/2, Fc5/6, Ft9/10, Tp7/8, Cz, C3/4, Cp1/2, Cp5/6, Pz, P3/4, P7/8, O1/2, Oz) and at the left and right mastoids. An electrode placed at the FCz position served as an online reference. EEG was re-referenced offline to the linked mastoids. Vertical eye movements were monitored with an electrode at the infraorbital ridge of the right eye. Electrode impedances were kept below 3 kΩ. An online notch filter at 50 Hz was used during the recording. EEG was re-referenced offline to the linked mastoids and data was band-pass filtered (0.1 Hz - 40 Hz) offline.

### Event-Related Potential (ERP) analysis

For each participant, we extracted the EEG epochs for each encoding odor + picture trial. Epochs had a duration of 10000 ms and included 6000 ms from odor onset and 4000 ms from the picture onset. EEG epochs were baseline corrected to the pre-stimulus interval (−100 to 0 ms). EEG trials with amplifier saturation, or trials with a shift exceeding 100 *µ*V/s were automatically rejected offline.

For subsequent analysis, including ERP and Neural Stability Analysis (see below), all the epochs were smoothed by averaging data via a moving window of 100 ms (excluding the baseline period) and then downsampled by a factor of 10. Statistical comparisons between conditions were made on the averaged ERP from all scalp electrodes.

### Neural Stability Analysis

We implemented a temporally resolved correlation analysis using Pearson coefficients [18]. The correlation analysis on EEG data was made at the individual level and to each time point separately and included spatial (i.e., scalp voltages from all the 29 electrodes) features of the resulting EEG single trials (Figure 4A). To examine the degree of EEG signal over time, we correlated, for each participant and at single trial level, the EEG patterns of activity elicited by pictures from each condition separately. This analysis was computed by randomly creating pairs of correlation analyses EEG trials from the same condition. For each participant, we created 200 permutations of possible unique pairings, and the final cross-correlation output resulted from averaging the point-to-point Fisher’s z scores correlation values across the 200 permutations.

### Nonparametric Cluster-Based Permutation Test

We implemented a cluster-based permutation test to account for neural stability differences between conditions [19]. This approach identifies clusters of significant points in the resulting 1D matrix in a data-driven manner and addresses the multiple-comparison problem by using a nonparametric statistical method based on cluster-level randomization testing to control the family-wise error rate. Statistics were computed for every time point, and the time points whose statistical values were larger than a threshold (p < 0.05, two-tail) were selected and clustered into connected sets based on adjacency points in the 1D matrix. The observed cluster-level statistics were calculated by taking the sum of the statistical values within a cluster. Condition labels were then permuted 1000 times to simulate the null hypothesis, and the maximum cluster statistic was chosen to construct a distribution of the cluster-level statistics under the null hypothesis. The nonparametric statistical test was obtained by calculating the proportion of randomized test statistics that exceeded the observed cluster-level statistics.

## Results

### Odor ratings

A repeated measures ANOVA unveiled variations in odor mean ratings (F(2,44) = 31.94, p < 0.01) as shown in Figure 2A. Ratings exceeding 4 were categorized as pleasant, while those below 4 were considered unpleasant. As expected, AIR consistently received a rating of 4, indicating its odorless nature (M = 4.01, SD = 0.06). CAR was slightly perceived as pleasant (M = 4.47, SD = 0.78) while BUT received an unpleasant rating (M = 2.99, SD = 0.70). Post-hoc paired student t-test comparisons further validated that CAR was rated as more pleasant than AIR (t(22) = 2.82, p = 0.01) and BUT (t(22) = 6.18, p < 0.01), while BUT was rated less pleasant than AIR (t(22) = −6.82, p < 0.01). These findings allow us to classify the different odor conditions into distinct hedonic categories, with CAR being positive, AIR being neutral, and BUT being negative.

**Figure 2.**
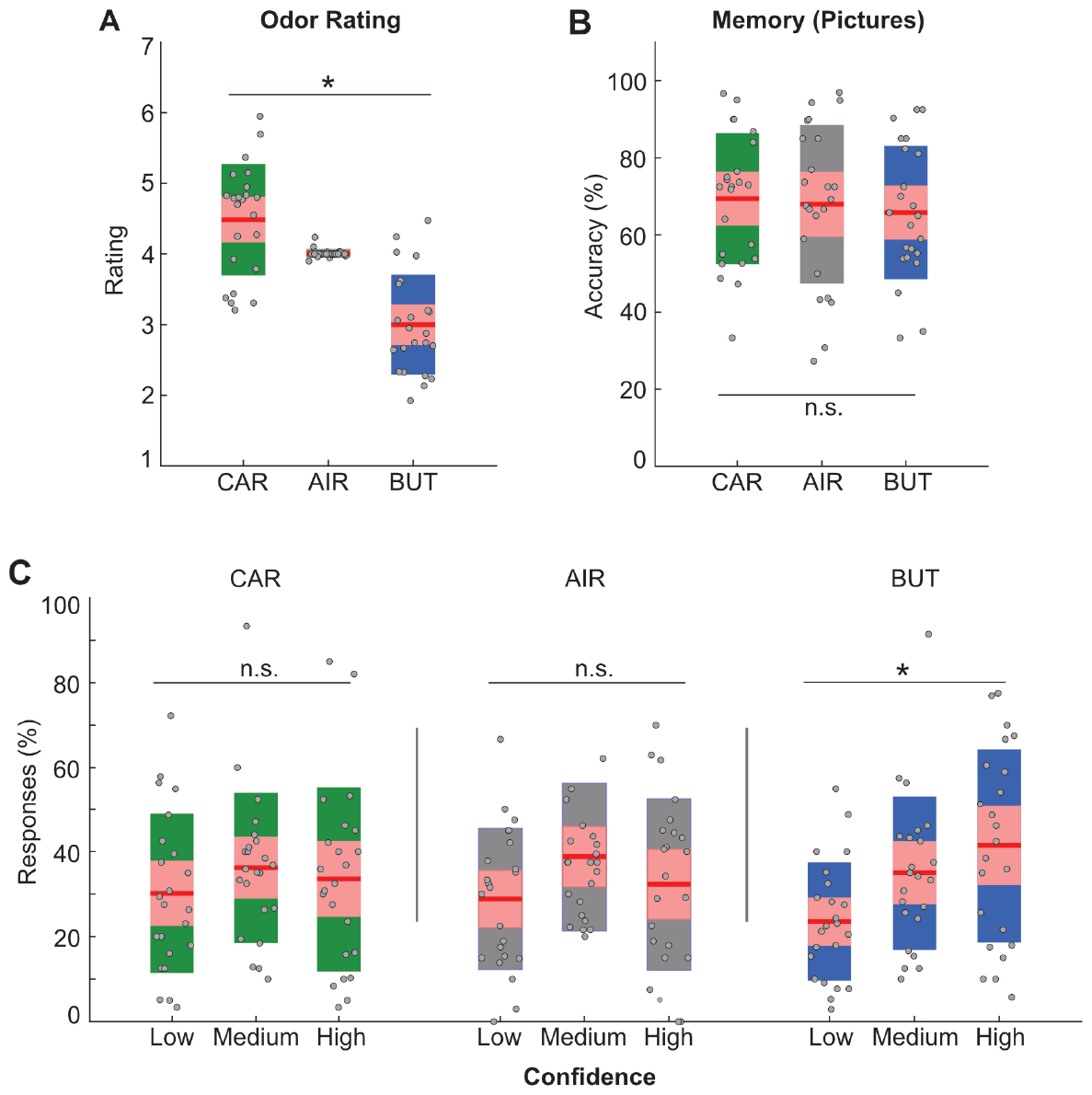
Behavioral results. (**A**) Odor ratings for L-Carvone (CAR), N-Butanol (BUT) and AIR odor conditions. (**B**) Percentage of remembered pictures associated to odor cue condition. (**C**) Percentage of confidence ratings associated to correct picture recognitions as a function their associated odor during encoding. ‘*’ indicates p ¡ 0.05. ‘n.s.’ indicates p ¿ 0.05.

### Memory performance for pictures

Odor presentation before the encoding of pictures did not play a role in the subsequent memory recognition task. Repeated measures ANOVA shows that there are not differences between odor conditions (F(2,44) = 0.62, p = 0.54) (Figure 2B).

A repeated measures ANOVA including Odor (CAR, AIR, and BUT) and confidence level (Low, Medium, and High) as within-subject factors, showed that participants’ correct recognition was linked to varying degrees of confidence depending on the type of odor (main effect of Confidence: F(2,44) = 1.28, p = 0.29; Odor x confidence interaction: F(4,88) = 4.48, p = 0.03). Further examination using a polynomial linear effect for this specific ANOVA revealed a linear increase in confidence ratings proportions (F(1,22) = 6.71, p = 0.02) (Figure 2C). To dissect this interaction effect, separate repeated measures ANOVAs were conducted for each Odor type. This analysis revealed that participants exhibited similar confidence levels for Hits associated with pictures linked to CAR (F(2,44) = 0.37, p = 0.69) and AIR (F(2,44) = 1.19, p = 0.31). However, confidence levels differed when pictures were encoded in connection with BUT (F(2,44) = 3.62, p = 0.03). Taken together, these results collectively suggest that the presence of odors could have impacted both the encoding and recall of associated pictures.

Conversely, participants showed to be accurate in correctly rejecting New items as Old during the memory recognition phase (M = 62.9%; SD = 19.8%). A repeated measures ANOVA, which included Confidence level (Low, Medium, and High) and Response type (Correct Rejection (CR) and False Alarm (FA)) as within subject factors, revealed a significant interaction between Confidence and Response type (F(2,44) = 8.36, p < 0.01), indicating that participants’ distribution of confidence varied between CR and FA. A series of post-hoc paired t-tests directly comparing these proportions between response types revealed that FA and CR were equally associated with High confidence ratings (t(22) = 1.64, p = 0.12). However, FA was linked to a greater number of low confidence responses compared to CR (Low: t(22) = −4.40, p < 0.01), while CRs exhibited a higher proportion of trials followed by intermediate confidence judgments (Medium: t(22) = 2.59, p = 0.02).

### Event-Related Potentials (ERPs)

The ERP analysis revealed that CAR and BUT odors elicited a greater EEG increase over centro-parietal electrodes compared to AIR conditions that persisted longer after their appearance (i.e., 0.5 s) (Figure 3). A cluster-based permutation analysis confirmed that CAR elicited a greater ERP than AIR from 657 to 2368 ms (cluster statistics: tsum = 138.51; tmax = 5.59, p = 0.04). A similar ERP increase was observed when comparing BUT and AIR, though this effect was longer in time (cluster statistics: tsum = 353.0; tmax = 6.78, p = 0.002; start/end = 657 – 4693 ms). We also found that BUT elicited a stronger ERP increase from 964 to 2588 ms than CAR (cluster statistics: tsum = 107.14; tmax = 3.88, p = 0.05). This analysis revealed that ERP changes between conditions were restricted to the period that comprised odor presentation upon the arrival of the picture, where similar ERP changes were found between conditions (all, p > 0.05).

**Figure 3.**
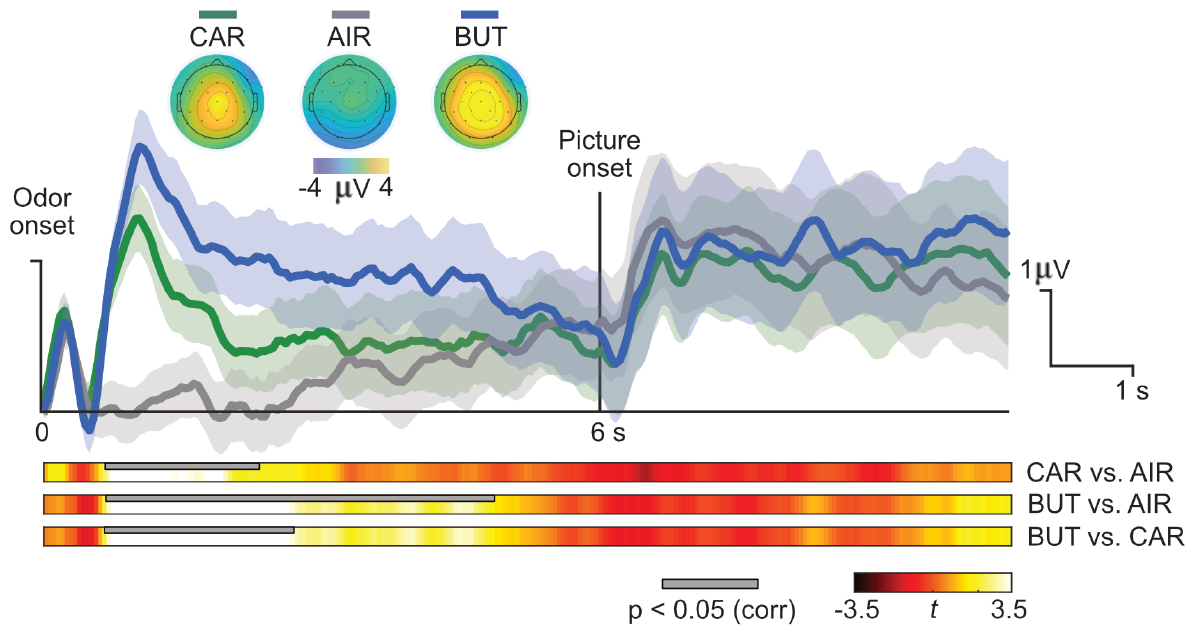
Event Related Potentials (ERPs). ERPs elicited by L-Carvone (CAR), N-Butanol (BUT) and AIR and the picture associated. Thick line display ERPs average over all 29 channels across participants. Point-to-point standard error of the mean (SEM) is depicted in shaded color. Point-to-point t values resulting from comparing paired conditions are displayed at the bottom. The thick grey line depicts the timing of the significant cluster between conditions (p < 0.05, cluster-based permutation test).

### Neural stability

We finally examined the extent to which EEG patterns of neural change elicited during odor + picture encoding elicited stable activity. This analysis revealed that the encoding of CAR and BUT elicited an increased pattern of neural activity that extended the odor exposure (i.e., 0.5s) and lasted for almost the encoding of the subsequent picture (Figure 4B). This effect was confirmed by the cluster-based permutation analysis, which revealed that CAR elicited greater neural stability than AIR during 517 to 3449 ms from odor onset (cluster statistics: tsum = 188.94; tmax = 4.20, p < 0.01, corrected). A similar effect was found for BUT, which elicited greater neural stability than AIR from 560 to 4556 ms (cluster statistics: tsum = 276.81; tmax = 3.81, p = 0.001, corrected). However, contrary to the ERP findings, CAR and BUT showed similar increase in neural stability during the entire EEG epoch of analysis (p > 0.05, corrected). Consistent with the ERP findings, no statistically significant differences in neural stability were found between conditions upon the appearance of the picture (all, p > 0.05, corrected). Please, note that the observed sudden increase in neural stability upon picture appearance is attributed to the similar ERP elicited by the encoding of a visual input.

**Figure 4.**
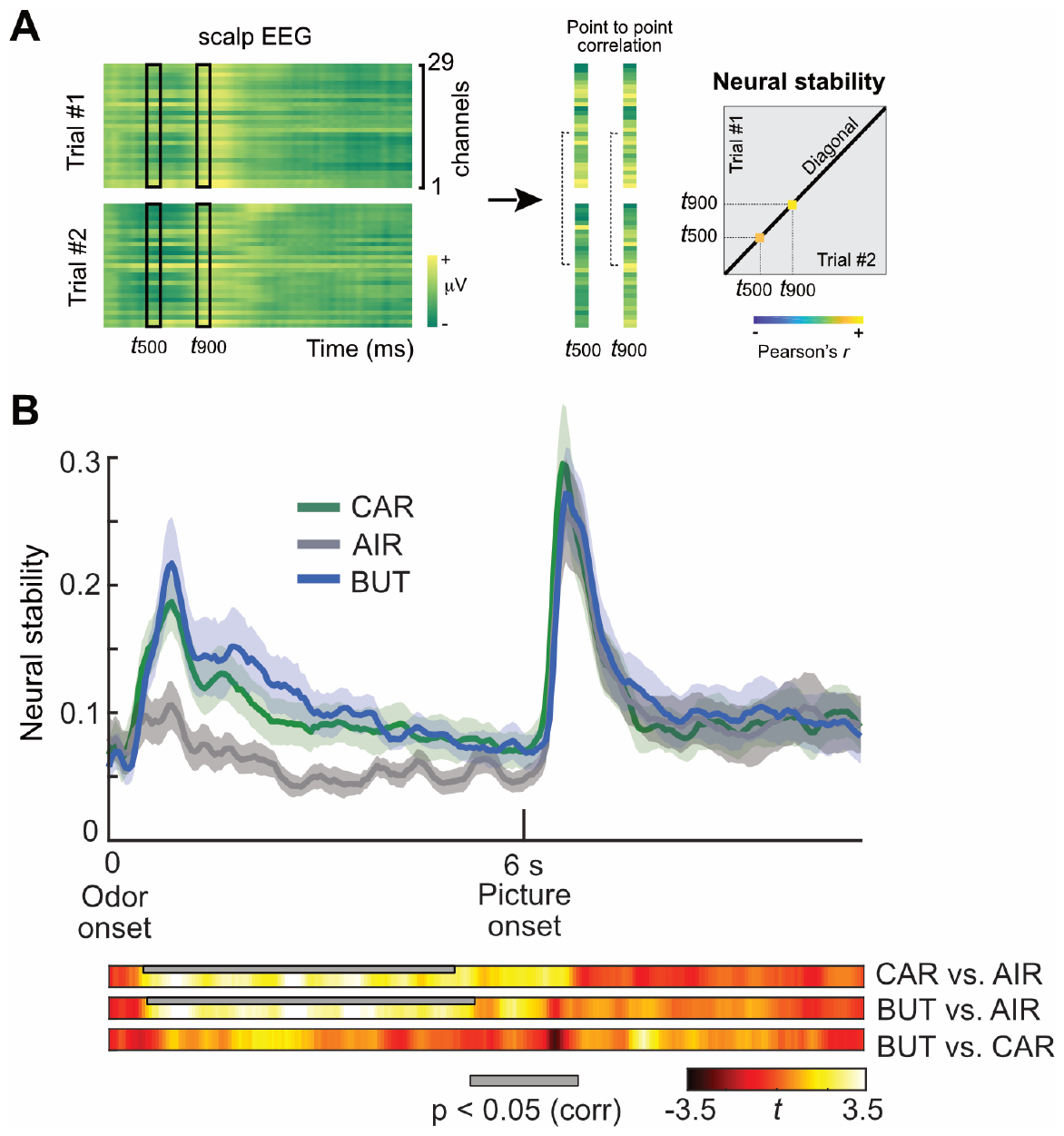
Neural stability. (**A**) Schematic representation of the analysis. A temporal cross-stimuli correlation matrix is generated from the EEG data for each participant. (**B**) Point-to-point participants’ average degree of neural stability for each odor and associated picture during encoding (thick line). The shaded area represents standard error (SEM) across subjects. Point-to-point t values resulting from comparing paired conditions are displayed at the bottom. The thick grey line depicts the timing of the significant cluster between conditions (p < 0.05, cluster-based permutation test).

## Discussion

In this study, we examined whether a brief exposure to odor cues triggered sustained neural activity during a 6 second delay period preceding the memory encoding of a picture. The combination of uni and multivariate ERP analytical approaches provided converging evidence that indeed, in line with nonhuman data [4, 5, 7], odor cues elicited, also in humans, a sustained neural activity that persisted beyond the odor exposure, enduring throughout the nearly 6-second delay period. Additionally, we found that the magnitude of the sustained ERP was higher for unpleasant than for pleasant odor cues. Finally, we found that participants remembered more confidently those pictures that were preceded by unpleasant than pleasant odor cues during their encoding, thereby indicating that brief odor cues influenced memory formation for temporally distant events.

Our findings align with the notion that persistent neural activity plays a crucial role in bridging events encoded separately in time and that it can been seen widespread over the brain [3]. Persistent neural activity has been observed in many cortical and subcortical regions of the human brain, notably in the prefrontal cortex, during the retention intervals of delayed response tasks [20]. While data from scalp EEG recordings cannot disambiguate the underlying brain sources of its activity, the combination of univariate and multivariate analysis on ERP data in the current study lends support to the possibility that distinct neural processes may take place during the delay period. Univariate ERP data revealed greater response amplitude for unpleasant than pleasant odor cues during the delay period whereas ERP signal was similarly stable for both pleasant and unpleasant odor cues throughout the delay period. Thus, while sustained neural processes show commonalities across olfactory stimuli regardless of hedonic dimensions, our results reveal that the magnitude of the brain response is sensitive to hedonic qualities.

The idea that sustained neural response dynamics were similar for pleasant and unpleasant odor cues but not present for neutral scents align with the idea that neural persistence detected from the scalp human EEG reflected more likely high-level processes derived from executive function, rather than sensory-specific representation. This idea would align with the long-held view that during the maintenance of sensory information in working memory relies in the coactivation of several neural coding systems, some that target sensory-specific memory maintenance, and other that enables the effective control of their maintenance upon interference [21]. Recent scalp ERP studies elicited by odor cues indicated that odor representation rapidly transforms in the brain reaching executive and semantic regions at around 800 ms from odor onset [11]. Our findings that odor cues elicited sustained ERPs that distinguished from neutral cues starting at around 650 ms would be in line with this rapid transformation dynamics in the brain view. A question that remains unresolved in our study relates to the sources of this ERP activity. The implementation of ERP approach to study odor elicited brain activity has shown to be sensitive to detect very early processing stages within the olfactory system, including the olfactory bulb [10], and expand larger brain areas associated with emotional, semantic processing [11]. Thus, a question for future studies would be to scrutinize whether sustained ERP activity and its impact in memory relates to the engagement of early vs late brain regions and how this interacted with the encoding of to-be-remembered material that relies on other sensory inputs, such as in the visual domain as in our study.

Our study revealed visual images that are preceded by unpleasant odors are better remembered than images preceded by pleasant or neutral odor cues. These findings are consistent with the idea that odor processing is intricately related to memory function [22] and is particularly sensible to aversive odors that could potentially be associated with a threat [10]. Indeed, previous human fMRI studies indicated that odors can modulate the neural processing visual inputs [23] and that odor-associated brain regions are reactivated during recall cued by visual cues [24]. However, two questions remained poorly investigated in the literature. First, while odor seems to be a powerful contextual parameter that influences memory, it is unclear if this modulation is brought by its subjective sense of hedonic characteristics. In fact, past studies showed that events that were encoded under pleasant odor contexts were better remembered in the future [24]. However, others indicated that memory benefit would be attributed by unpleasant odor contexts [25]. Second, while it is generally assumed that the benefit of odors in memory can be attributed to their link to emotional processing [26], other aspects related to how memories are formed can also play an important role. For example, research has indicated that odors impact in memory via activation of the amygdala [25], which is a powerful region of memory enhancement [27, 28]. However, other studies indicated that the involvement of hippocampus in odor-mediated memory enhancement is more relevant [24], as it is through hippocampal engagement that disparate encoding inputs from different modalities can become bound in a relational memory structure [29]. Thus, while these two views can be complementary, it would be important that future studies help disambiguate the exact contribution of each other. While the current study cannot directly speak about the involvement of the hippocampus and/or amygdala during encoding, our findings that ERP signal differ during the delay period after odor encoding may fit better to the idea that the odor-induced memory modulation of image pictures to be attributed to associative processing rather than transferred emotional processing during the encoding of the picture. We reasoned that if the latter was the case, we would have observed changes in the ERP signals, in line with previous data showing that ERPs are sensitive to inputs with emotional content elicited by unpleasant odors [11, 30, 31].

In conclusion, our study establishes that brief exposure to odor cues induces sustained neural activity in humans, echoing patterns observed in nonhuman subjects. The greater amplitude of sustained ERPs for unpleasant odors underscores the distinct neural responses associated with different odor valences. Furthermore, the observed impact on memory recall emphasizes the influential role of brief odor cues in shaping and enhancing the encoding of temporally distant events.

## Author contributions

J.T., J.M.P., and L.F. designed the experiment. J.T., M.M. and J.V. conducted the experiment. J.T., M.M., J.M.P. and L.F. analyzed the data. D.P. and J.V. contributed to the chemical advisory as well as in the acquisition and dilution of the olfactory stimuli. J.T. and L.F. wrote the original draft. J.M.P. reviewed and edited the manuscript. All authors have read and agreed to the published version of the manuscript.

## Acknowledgments

We thank Xiongbo Wu and Josué García-Arch for their help during the data acquisition and the analyses at early stages of this study. This work was supported by the Spanish Ministerio de Ciencia, Innovación y Universidades, which is part of Agencia Estatal de Investigación (AEI), through the project PID2022 - 140426NB - I00 (funded by MCIN/ AEI /10.13039/501100011033/ and FEDER a way to make Europe), to L.F. and from Puig S.L. We thank CERCA Programme/Generalitat de Catalunya for institutional support. J.T. is supported by a grant from Puig S.A.

## References

1. Joaquin M. Fuster. Memory in the cerebral cortex: an empirical approach to neural networks in the human and nonhuman primate. MIT Press, Cambridge, Mass, 1995.

2. P.S Goldman-Rakic. Cellular basis of working memory. Neuron, 14(3):477–485, March 1995.

3. Guy Major and David Tank. Persistent neural activity: prevalence and mechanisms. Current Opinion in Neurobiology, 14(6):675–684, December 2004.

4. Michael Andrew Patterson, Samuel Lagier, and Alan Carleton. Odor representations in the olfactory bulb evolve after the first breath and persist as an odor afterimage. Proceedings of the National Academy of Sciences, 110(35), August 2013.

5. Dana Shani Galili, Alja Lüdke, C. Giovanni Galizia, Paul Szyszka, and Hiromu Tanimoto. Olfactory Trace Conditioning in Drosophila. The Journal of Neuroscience, 31(20):7240–7248, May 2011.

6. Heather Carlson, Joana Leitão, Sylvain Delplanque, Isabelle Cayeux, David Sander, and Patrik Vuilleumier. Sustained effects of pleasant and unpleasant smells on resting state brain activity. Cortex, 132:386–403, November 2020.

7. Xiaoxing Zhang, Wenjun Yan, Wenliang Wang, Hongmei Fan, Ruiqing Hou, Yulei Chen, Zhaoqin Chen, Chaofan Ge, Shumin Duan, Albert Compte, and Chengyu T Li. Active information maintenance in working memory by a sensory cortex. eLife, 8:e43191, jun 2019.

8. Leslie M. Kay. Circuit Oscillations in Odor Perception and Memory. In Progress in Brain Research, volume 208, pages 223–251. Elsevier, 2014.

9. Ko Kobayakawa, Reiko Kobayakawa, Hideyuki Matsumoto, Yuichiro Oka, Takeshi Imai, Masahito Ikawa, Masaru Okabe, Toshio Ikeda, Shigeyoshi Itohara, Takefumi Kikusui, Kensaku Mori, and Hitoshi Sakano. Innate versus learned odour processing in the mouse olfactory bulb. Nature, 450(7169):503–508, November 2007.

10. Behzad Iravani, Martin Schaefer, Donald A. Wilson, Artin Arshamian, and Johan N. Lundström. The human olfactory bulb processes odor valence representation and cues motor avoidance behavior. Proceedings of the National Academy of Sciences, 118(42):e2101209118, October 2021.

11. Mugihiko Kato, Toshiki Okumura, Yasuhiro Tsubo, Junya Honda, Masashi Sugiyama, Kazushige Touhara, and Masako Okamoto. Spatiotemporal dynamics of odor representations in the human brain revealed by EEG decoding. Proceedings of the National Academy of Sciences, 119(21):e2114966119, May 2022.

12. Mäellie Midroit, Laura Chalençon, Nicolas Renier, Adrianna Milton, Marc Thevenet, Jöelle Sacquet, Marine Breton, Jérémy Forest, Norbert Noury, Marion Richard, Olivier Raineteau, Camille Ferdenzi, Arnaud Fournel, Daniel W. Wesson, Moustafa Bensafi, Anne Didier, and Nathalie Mandairon. Neural processing of the reward value of pleasant odorants. Current Biology, 31(8):1592–1605.e9, April 2021.

13. C. C. Licon, C. Manesse, M. Dantec, A. Fournel, and M. Bensafi. Pleasantness and trigeminal sensations as salient dimensions in organizing the semantic and physiological spaces of odors. Scientific Reports, 8(1):8444, May 2018.

14. Howard R. Moskowitz, Andrew Dravnieks, and Clifford Gerbers. Odor intensity and pleasantness of butanol. Journal of Experimental Psychology, 103(2):216–223, August 1974.

15. Javiera P. Oyarzún, Pau A. Packard, Ruth De Diego-Balaguer, and Lluis Fuentemilla. Motivated encoding selectively promotes memory for future inconsequential semantically-related events. Neurobiology of Learning and Memory, 133:1–6, September 2016.

16. Alex Holkner and Pyglet Developers. Pyglet: A cross-platform windowing and multimedia library for Python. https://pyglet.org/, 2021.

17. Jonathan Peirce, Jeremy R. Gray, Sol Simpson, Michael MacAskill, Richard Höchenberger, Hiroyuki Sogo, Erik Kastman, and Jonas Kristoffer Lindeløv. PsychoPy2: Experiments in behavior made easy. Behavior Research Methods, 51(1):195–203, February 2019.

18. Xiongbo Wu and Llúis Fuentemilla. Distinct encoding and post-encoding representational formats contribute to episodic sequence memory formation. Cerebral Cortex, 33(13):8534–8545, June 2023.

19. Eric Maris and Robert Oostenveld. Nonparametric statistical testing of EEG- and MEG-data. Journal of Neuroscience Methods, 164(1):177–190, August 2007.

20. Clayton E. Curtis and Mark D’Esposito. Persistent activity in the prefrontal cortex during working memory. Trends in Cognitive Sciences, 7(9):415–423, September 2003.

21. Mark D’Esposito and Bradley R. Postle. The Cognitive Neuroscience of Working Memory. Annual Review of Psychology, 66(1):115–142, January 2015.

22. Rachel S. Herz and Trygg Engen. Odor memory: Review and analysis. Psychonomic Bulletin & Review, 3(3):300–313, September 1996.

23. Jay A Gottfried and Raymond J Dolan. The Nose Smells What the Eye Sees. Neuron, 39(2):375–386, July 2003.

24. Jay A Gottfried, Adam P.R Smith, Michael D Rugg, and Raymond J Dolan. Remembrance of Odors Past. Neuron, 42(4):687–695, May 2004.

25. Stephan B. Hamann, Timothy D. Ely, Scott T. Grafton, and Clinton D. Kilts. Amygdala activity related to enhanced memory for pleasant and aversive stimuli. Nature Neuroscience, 2(3):289–293, March 1999.

26. Rachel Herz. The Role of Odor-Evoked Memory in Psychological and Physiological Health. Brain Sciences, 6(3):22, July 2016.

27. R Adolphs, L Cahill, R Schul, and R Babinsky. Impaired declarative memory for emotional material following bilateral amygdala damage in humans. Learn Mem, 4, 291–300, 1997.

28. Larry Cahill, Ralf Babinsky, Hans J. Markowitsch, and James L. McGaugh. The amygdala and emotional memory. Nature, 377(6547):295–296, September 1995.

29. Neal J. Cohen, Russell A. Poldrack, and Howard Eichenbaum. Memory for Items and Memory for Relations in the Procedural/Declarative Memory Framework. Memory, 5(1-2):131–178, January 1997.

30. Thomas Hummel and Gerd Kobal. Differences in human evoked potentials related to olfactory or trigeminal chemosensory activation. Electroencephalography and Clinical Neurophysiology/Evoked Potentials Section, 84(1):84–89, January 1992.

31. J. N. Lundstrom, S. Seven, M. J. Olsson, B. Schaal, and T. Hummel. Olfactory Event-Related Potentials Reflect Individual Differences in Odor Valence Perception. Chemical Senses, 31(8):705–711, October 2006.

